# Thrombocytopenia Microcephaly Syndrome - a novel phenotype associated with *ACTB* mutations

**DOI:** 10.1101/303909

**Authors:** Sharissa L. Latham, Nadja Ehmke, Patrick Y.A. Reinke, Manuel H. Taft, Michael J. Lyons, Michael J Friez, Jennifer A. Lee, Ramona Hecker, Michael C. Frühwald, Kerstin Becker, Teresa M. Neuhann, Denise Horn, Evelin Schrock, Katharina Sarnow, Konrad Grützmann, Luzie Gawehn, Barbara Klink, Andreas Rump, Christine Chaponnier, Ralf Knöfler, Dietmar J. Manstein, Natalia Di Donato

**Affiliations:** Institute for Biophysical Chemistry, Hannover Medical School, Hannover, Germany.; Institute of Medical and Human Genetics, Charité-Universitätsmedizin Berlin, Berlin, Germany.; Berlin Institute of Health, Berlin, Germany.; Division for Structural Biochemistry, Hannover Medical School, Hannover, Germany.; Greenwood Genetic Center, Greenwood, South Carolina, USA.; Institute for Clinical Chemistry and Laboratory Medicine, Medical Faculty of TU Dresden, Dresden, Germany.; Swabian Children’s Cancer Center, Children’s Hospital Augsburg, Augsburg, Germany.; Medical Genetics Center, Munich, Germany.; Institute for Clinical Genetics, TU Dresden, Dresden, Germany.; Core Unit for Molecular Tumor Diagnostics, National Center for Tumor Diseases Dresden, Dresden, Germany.; Department of Pathology-Immunology, Faculty of Medicine, University of Geneva, Geneva, Switzerland.; Department of Paediatric Haemostaseology, Medical Faculty of TU Dresden, Dresden, Germany

## Abstract

Until recently missense germ-line mutations in *ACTB*, encoding the ubiquitously expressed β-cytoplasmic actin (CYA), were exclusively associated with Baraitser-Winter Cerebrofrontofacial syndrome (BWCFF), a complex developmental disorder^1,2^. Here, we report six patients with previously undescribed heterozygous variants clustered in the 3’-coding region of *ACTB*. These patients present with clinical features different from BWCFF, including thrombocytopenia, microcephaly, and mild developmental disability. Patient derived cells are morphologically and functionally distinct from controls. Assessment of cytoskeletal constituents identified a discrete filament population altered in these cells, which comprises force generating and transmitting actin binding proteins (ABP) known to be associated with thrombocytopenia^3–8^. *In silico* modelling and molecular dynamics (MD)-simulations support altered interactions between these ABP and mutant β-CYA. Our results describe a new clinical syndrome associated with *ACTB* mutations with a distinct genotype-phenotype correlation, identify a cytoskeletal protein interaction network crucial for thrombopoiesis, and provide support for the hypomorphic nature of these actinopathy mutations.

## Main Text

The human genome encodes six actin isoforms, produced in a time- and tissue-specific manner during development. Isoforms are classified according to their relative isoelectric focusing mobility and enrichment in striated muscle (α-cardiac and α-skeletal actins), smooth muscle (α-smooth and γ-smooth actins (SMA)), and nonmuscle (β-cytoplasmic and γ-cytoplasmic actins (CYA)) tissues^9^. All are highly conserved throughout evolution (>93% identity) and are associated with disease-causing mutations^10,11^. Mutations in *ACTB* and *ACTG1*, encoding the ubiquitously expressed CYA isoforms, are associated with a broad spectrum of clinical phenotypes. Heterozygous germ-line mutations in both genes have been associated with BWCFF, a well-defined syndrome with recognizable facial features, developmental disability, neuronal migration defects, hearing loss, ocular colobomas, heart and renal defects, and progressive muscle wasting^2^. The consequences of *ACTG1* germ-line mutations are less pleiotropic. They have been linked to isolated non-syndromic hearing loss^12^. Moreover, two recent reports have identified *ACTB* post-zygotic mutations and *ACTB* haploinsufficiency due to 7p22.1 microdeletion in patients with Becker’s Nevus Syndrome^13^ and a pleotropic developmental disorder^14^.

Through whole-exome or large gene-panel sequencing, we identified 6 patients (P1-6) from 4 unrelated families carrying *de novo* or co-segregating heterozygous *ACTB* variants (Table 1, Supplementary Table 1). These mutations cluster in the 3’ region of the gene (Fig. 1), within exons 5 and 6, encoding residues 313–368 of β-CYA (Supplementary Fig. 1). None of these patients showed features associated with BWCFF, for which missense mutations occur within exons 2–4. Transitory or permanent low blood thrombocyte count is common to all patients. Thrombocyte anisotropy is observed in peripheral blood smears and the fraction of immature thrombocytes is elevated (37–45%; reference range: 1–6%). Bone marrow examination showed an increased megakaryocyte count in P5 (Supplementary Fig. 2). Additionally, all mutations are associated with microcephaly. Intellectual function ranges from mild developmental disability to normal intellect. Facial dysmorphism, if observed, appears to be distinct from the typical facial features of BWCFF patients (Table 1, Supplementary Fig. 3). Given the distinct genotype-phenotype correlation, we name this actinopathy *ACTB*-associated <u>T</u>hrombocytopenia <u>M</u>icrocephaly <u>S</u>yndrome (*ACTB*-TMS).

**Figure 1.**
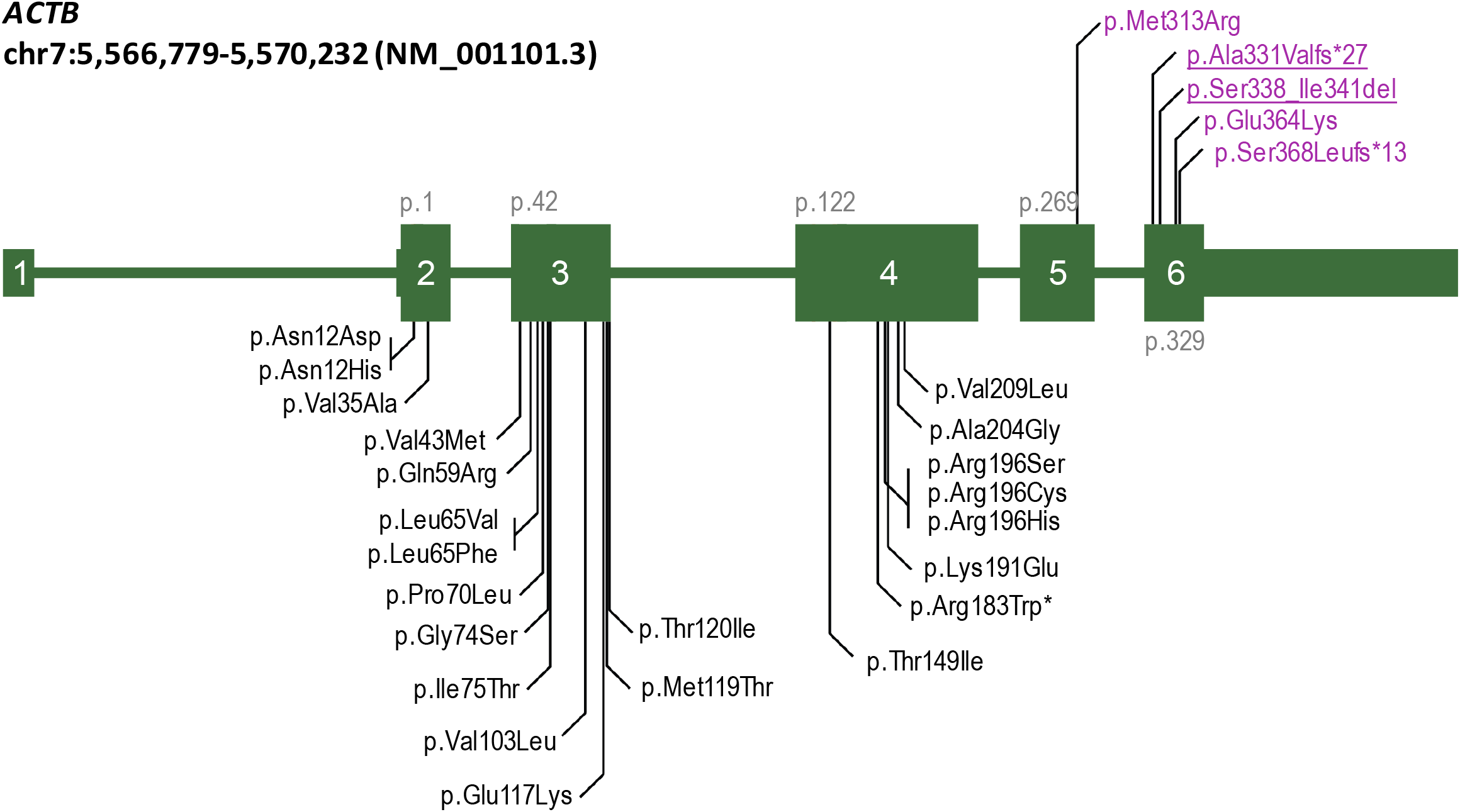
Schematic representation of *ACTB*-TMS mutations. Genomic coordinates refer to GRCh37/hg19 genome assembly (modified from Verloes *et al*^2^); exons are numbered, and coding exons are indicated by large boxes; BWCFF associated mutations from the literature and unpublished mutations from the own cohort are shown below the gene model; *ACTB*-TMS are highlighted in magenta above the gene model; * indicates a specific amino acid change associated with progressive dystonia^61^; mutations studied on the cellular level in the present work are underlined.

**Table 1.** Clinical overview of *ACTB*-TMS mutations and patient cohort.

| **A. Summary of *ACTB*-TMS mutations.** |  |  |  |
| --- | --- | --- | --- |
| **Patient** | **Genomic position (hg19)** | **cDNA (NM_001101.3)** | **Amino acid change** |
| 1<sup>*</sup>, 2 | chr7:g.[5567681A>C] | c.938T>G | p.(Met313Arg) |
| 3<sup>#</sup>, 4 | chr7:g.[5567499_5567515del] | c.992_1008del | p.(Ala331Valfs*27) |
| 5<sup>*,#,\$</sup> | chr7:g.[5567484_5567495del] | c.1012_1023del | p.(Ser338_Ile341del) |
| 6 | chr7:g.[5567406dup] | c.1101dup | p.(Ser368Leufs*13) |
| N<sup>‡</sup> | chr7:g.[5567417C>T] | c.1090G>A | p.(Glu364Lys) |
| **B. Clinical overview of individuals with *ACTB*-TMS.** |  |  |  |
| **Features** |  |  | **Number / total tested** |
| **Microcephaly** |  |  | 5 / 6 |
| **Facial minor anomalies** |  |  |  |
| Flared and straight eyebrows |  |  | 6/6 |
| Synophrys |  |  | 3/6 |
| Epicanthic folds |  |  | 4/6 |
| Broad nasal bridge and tip |  |  | 6/6 |
| Thin upper vermilion border |  |  | 4/6 |
| **Hematological anomalies** |  |  |  |
| Thrombocytopenia |  |  | 6/6 |
| Anisotropy with enlarged platelets |  |  | 5/5<sup>(P2-6)</sup> |
| High immature platelet fraction |  |  | 2/2<sup>(P2, P5)</sup> |
| Increased eosinophilic granulocytes |  |  | 2/6 |
| **Mild developmental disability** |  |  | 5/6 |
| Features observed in single individuals: heart defect, incomplete cleft lip, periventricular nodular heterotopias, monocytosis, and leukocytosis. |  |  |  |

^*^Access to peripheral blood smears for thrombocyte immunofluorescence

^#^Access to primary dermal fibroblasts for cell-based assays and RNA-sequencing analysis

^§^Considered to be the most severely affected patient in this *ACTB*-TMS cohort based on severe microcephaly, prominent facial features and hematological anomalies

^‡^Patient described by Nunoi et al. and the only case where mutant β-CYA has been biochemically characterised^36^

Different from *ACTB*-TMS patients, BWCFF patients have not been reported to display abnormal blood counts^2^. This excludes possible associations with hematological malignancies^15^. One patient with a missense mutation in exon 6 of *ACTB* displaying moderate intellectual disability and thrombocytopenia, in addition to abnormal white blood cell counts and recurrent infections, was previously reported^16^ (Supplementary Table 1, Patient N). As no cranial measurements were documented by Nunoi *et al*., we suggest that this patient shares typical *ACTB*-TMS symptoms. P5, who presented with the most severe clinical symptoms in our cohort, displays partially overlapping white blood cell abnormalities as PN but without leukocytopenia and recurrent infections.

To assess how *ACTB*-TMS variants affect cytoskeletal organization and cellular function, primary dermal fibroblasts were harvested from P3 and P5. Prominent morphological differences are observed between control and *ACTB*-TMS fibroblasts (Fig. 2a). Patient cells cover significantly less substrate surface area than the healthy control (Fig. 2b) and have ~25% reduced total cell volume (Fig. 2c). A distinct feature of P5 fibroblasts is their organization in tightly packed clusters (Fig. 2a, arrow).

**Figure 2.**
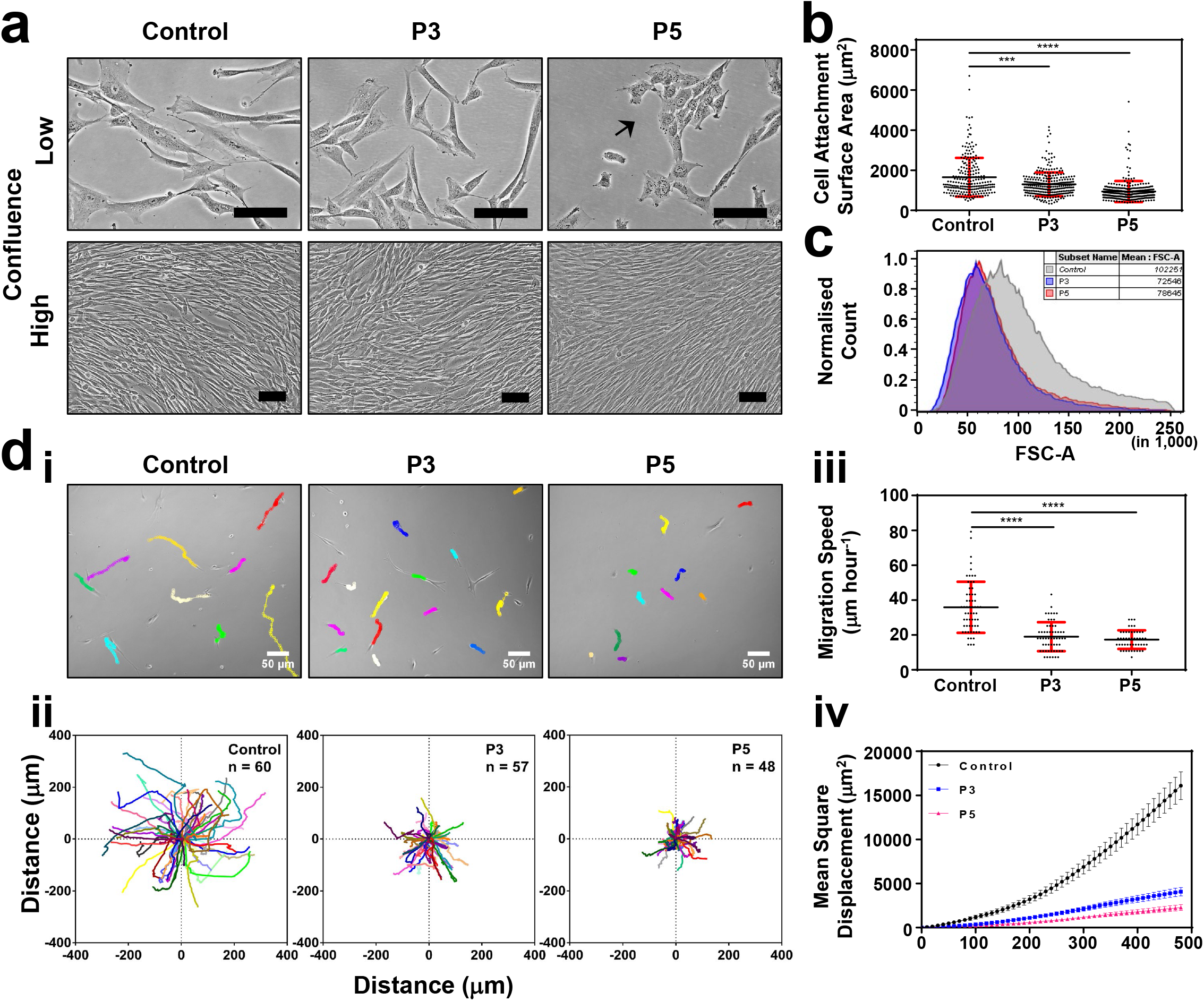
The total volume, cell attachment surface area and migratory capacity of *ACTB*-TMS dermal fibroblasts is reduced compared to healthy control fibroblasts. (**a**) Micrographs of control, P3 (p.Ala331Valfs*27) and P5 (p.Ser338_Ile341del) primary dermal fibroblasts at low (top) and high (bottom) confluence. *ACTB*-TMS cells are distinctly smaller and P5 cells aggregate with one another. Scale bars are 100 μm; (**b**) dot plot of cell attachment surface area shows reduced coverage by *ACTB*-TMS cells (mean ± SD, **** = P<0.0001); (**c**) Forward Scatter Area (FSC-A) vs normalized cell count flow cytometry analysis shows a reduction in the total volume of *ACTB*-TMS cells (P3: blue, P5: red) compared to controls (grey); (**d**) migration of control and *ACTB*-TMS primary fibroblasts, demonstrating reduced migratory capacity for patient cells; (**i**) representative tracks at 8 h; (**ii**) trajectories of all tracks for control (left), P3 (mid) and P5 (right) fibroblast movement from 0–8 h; (**iii**) dot plots of cell migration speed represented in μm per minute (mean ± SD; **** = P<0.0001); (**iv**) mean square displacement analysis of control (black), P3 (blue) and P5 (red) fibroblasts show that unlike controls, *ACTB*-TMS cells do not display persistent migration.

Migration assays show that this occurs as cell-cell interactions cannot be disrupted (Supplementary Fig. 4). Examination of cell trajectory, migration speed and persistence show that ACTB-TMS cells exhibit limited migratory capacity compared with control cells. No significant differences in cell proliferation rate are observed between the cultures (Supplementary Fig. 4).

Despite differing by only four N-terminal residues, β-CYA and γ-CYA are divergent in their cellular localizations and functions. In the cytoplasm, γ-CYA localizes to sub-membranous networks and promotes cell migration, β-CYA is implicated in contraction and enriches in stress fibers and cellcell contacts^17–19^. In control and patient fibroblasts, we observe discrete lateral and axial isoactin segregation (Fig. 3a, Supplementary Fig. 5). As previously reported^19^, β-CYA is enriched in sub-nuclear filaments at the cells’ basal plane, whilst γ-CYA is enriched at the cell periphery. These β-CYA sub-nuclear filaments are consistent between healthy control and BWCFF p.Thr120Ile control fibroblasts (P1 in ^20^), but are phenotypically distinct in ACTB-TMS cells (Fig. 3b). Here, filaments are bundled into thick fibers and γ-CYA levels are reduced. Despite enrichment of β-CYA at this locality, total protein analysis shows reduced β-CYA levels and a compensatory increase in γ-CYA, whilst total actin is unchanged (Fig. 3c).

**Figure 3.**
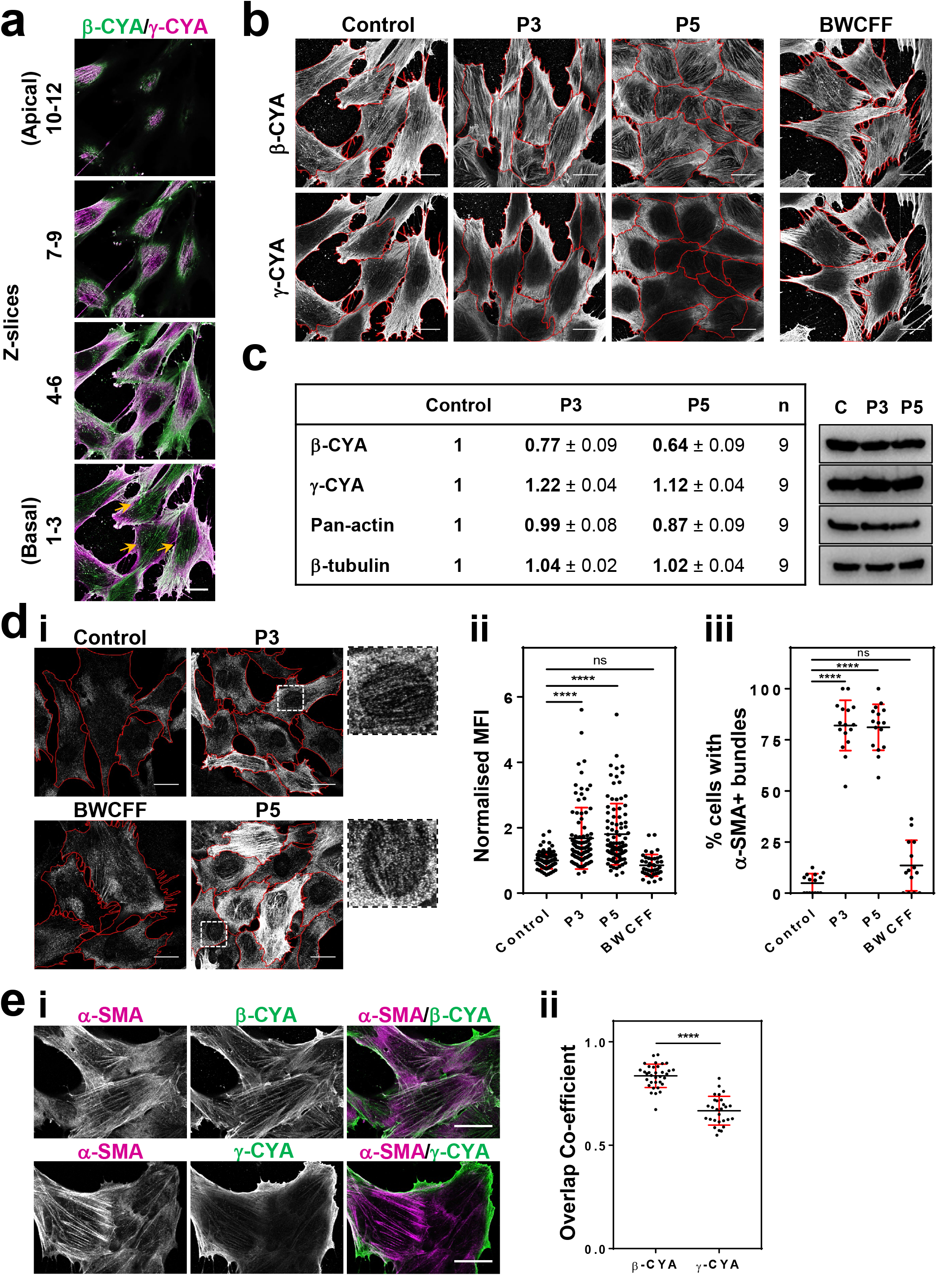
α-SMA compensates for reduced β-CYA levels in ACTB-TMS patient primary fibroblasts, where it is preferentially incorporated into sub-nuclear β-CYA bundles. (**a**) β-CYA (green) and γ-CYA (magenta) distribution in healthy control fibroblasts shows discrete lateral and axial isoform segregation. Maximum intensity projections for indicated z-stack slices are given; (**b**) maximum intensity projections of z-stack slices 1–2 are given for β-CYA (top row) and γ-CYA (bottom row) in co-stained healthy control (left), P3 (p.Ala331Valfs*27) (mid left), P5 (p.Ser338_Ile341del) (mid right) and BWCFF (p.Thr120Leu) control (right) fibroblasts. β-CYA is organized into thick, sub-nuclear bundles in P3 and P5; (**c**) Western blot analysis of control and patient fibroblasts showing reduced β-CYA levels and elevated α-SMA levels are associated with ACTB-TMS mutations. Protein levels are represented relative to the control (mean ± SE); (**d**) (**i**) immunofluorescence staining of α-SMA shows increased expression in P3 and P5 fibroblasts (inserts on right); (**ii**) mean fluorescence intensity (MFI) values of individual cells relative to the control average are given (± SD); (**iii**) quantification of the percentage of cells per field where α-SMA is incorporated into basal sub-nuclear filaments (n=12–14); (**e**) (**i**) P5 cells co-stained for α-SMA (magenta) with β-CYA (top, green) or γ-CYA (bottom, green), showing (**ii**) greater overlaps of α-SMA with β-CYA within the basal plane of individual cells. All scale bars represent 20 μm and red markings show cell boundaries. In all cases, **** = P<0.0001 and ns = not significant.

Although typical of vascular smooth muscle tissue, α-SMA is also produced by dermal fibroblasts that undergo myofibroblast differentiation to promote final wound contraction during healing processes^21^. Higher levels of α-SMA are therefore correlated with increased contractile activity, enhanced cell-cell contacts and reduced migratory capacity^22–24^, consistent with the phenotype of *ACTB*-TMS fibroblasts. As α-SMA is upregulated in *ACTB^−/−^* mouse embryonic fibroblasts^24^, we assessed its expression in *ACTB*-TMS fibroblasts. Strong signals are observed in both patient cultures by Western blotting (Supplementary Fig. 5). Whilst great variability in α-SMA levels is observed between neighboring cells by immunofluorescence microscopy, the average mean fluorescence intensity (MFI) is significantly increased in P3 and P5 cells compared to healthy and BWCFF controls (Fig. 3d). α-SMA localizes to basal sub-nuclear filaments in ~80% of *ACTB*-TMS cells (Fig. 3d.i and iii). Colocalization analysis shows a higher degree of overlap between α-SMA and β-CYA compared to γ-CYA, indicating preferential incorporation of α-SMA into these β-CYA bundles (Fig. 3e, Supplementary Fig. 5).

Whole-transcriptome (RNA-Seq) analysis was performed to identify genes, specifically those encoding ABPs, significantly deregulated in P3 and P5 primary fibroblasts compared to healthy control primary fibroblasts (Supplementary Tables 2–4). Of the 124 ABP-encoding RNAs detected at high levels in control or patient cells (Fig. 4a.i, Supplementary Tables 5–6), 34 are significantly upregulated in both patients and in a manner consistent with clinical severity (log2 fold change (FC) P5>P3, Fig. 4a.ii). Amongst these, five candidate genes encode ABPs implicated in thrombocytopenia, the most consistent phenotype in *ACTB*-TMS. These genes encode for α-actinin-1^34^, nonmuscle myosin-2A (NM-2A)^5,6^, diaphanous formin-1 (Diaph1)^25^, filamin A^7,8^ and tropomyosin (Tpm)4.2^26^. Total protein analysis shows that Diaph1 is unchanged between control and *ACTB*-TMS samples, whilst all other gene products are increased (Fig. 4b). By immunofluorescence, we demonstrate enrichment of α-actinin-1 and NM-2A, and the recruitment of filamin A, into basal β-CYA/α-SMA bundles in both P3 and P5 fibroblasts (Fig. 4c). An antibody recognizing Tpm4.1/4.2 isoforms does not localize to these structures (Supplementary Fig. 6).

**Figure 4.**
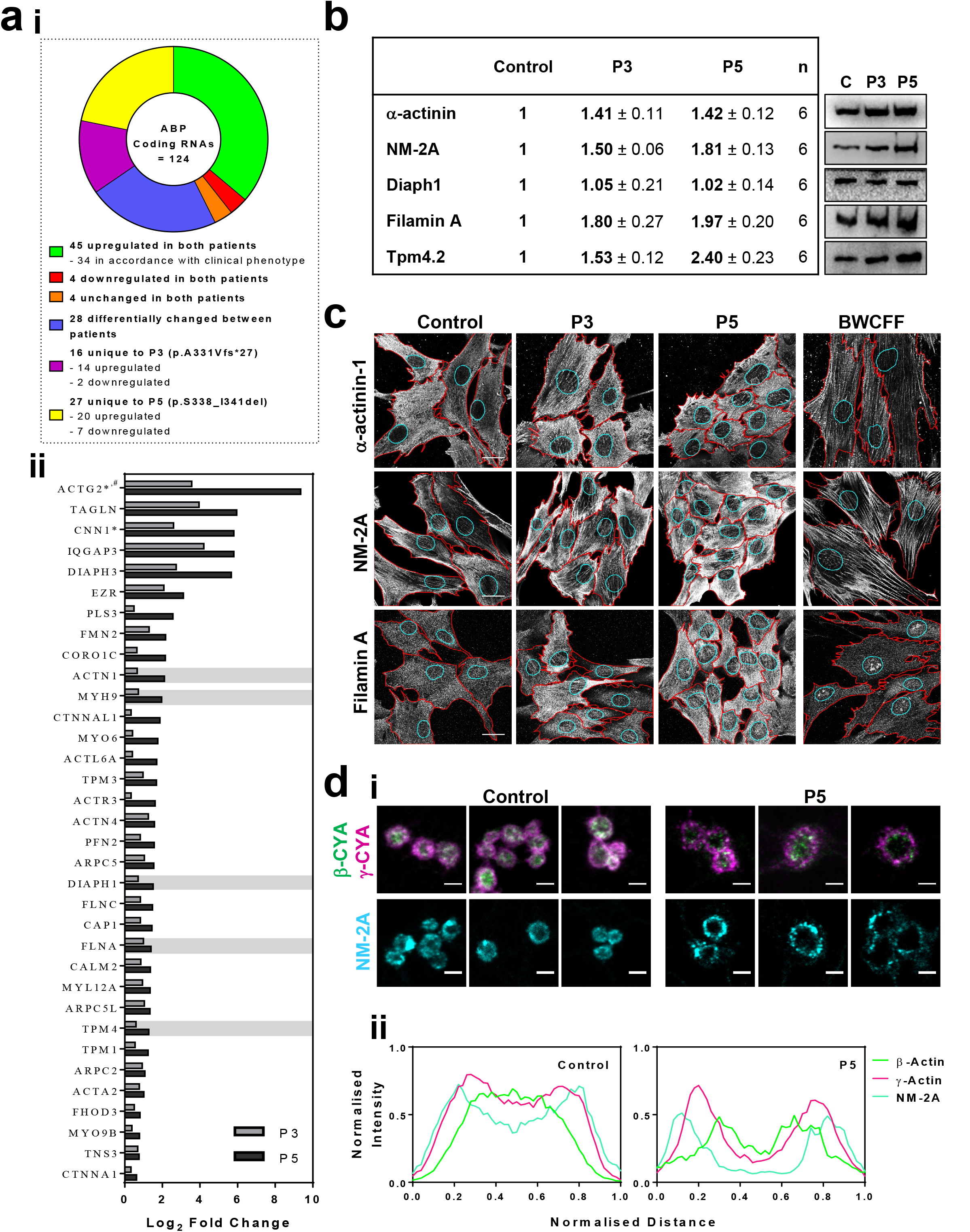
Aberrant recruitment and distribution of thrombocytopenia-associated ABPs, NM-2A, α-actinin and filamin A in *ACTB*-TMS primary fibroblasts and peripheral blood thrombocytes. (**a**) The total RNA content of control, P3 (p.Ala331Valfs*27) and P5 (p.Ser338_Ile341del) primary fibroblasts was examined by whole-transcriptome analysis; (**i**) a subset of 124 ABP genes were detected with >300 reads in control or patient fibroblasts. Of these, 120 were significantly deregulated in one or both patients compared to the control; (**ii**) Of the 34 genes upregulated in both patients in accordance with clinical severity (log_2_ FC of P5>P3), 5 genes with known thrombocytopenia-associated mutations were identified (grey shading). * = genes with < 20 reads in all 3 control measurements, # = no protein detected by Western Blot; (**b**) Western Blot analysis of control vs. patient fibroblasts shows increased expression of all candidates, except for Diaph1. Data are given as FC relative to the control (mean ± SE); (**c**) Maximum intensity projections of z-stack slices 1–2 are given for α-actinin (top row) NM-2A (middle row), filamin A (bottom row), showing recruitment of these ABP to the basal, subnuclear filament population in P3 and P5 samples, and not in healthy or BWCFF (p.Thr120Leu) controls. Cell boundaries are red, nuclear boundaries are cyan and scale bars represent 10 μm; (**d**) (**i**) Immunofluorescence staining of control and P5 peripheral blood smears labelled for β-CYA (green), γ-CYA (magenta) and NM-2A (cyan). Scale bars represent 2 μm; (**ii**) Line scan analysis (normalized data combined from 10 events) shows reduced and discontinuous staining in patient thrombocytes.

Cellular contractile force is generated through ATP-dependent interactions of NM-2 isoforms and filamentous actin^27,28^. Crosslinkers such as α-actinin stabilize and regulate these interactions, and in combination with the membrane anchoring filamin A, transmit this force to the plasma membrane at focal adhesions, cell-cell junctions, and through interactions with a subset of transmembrane proteins^29,30^. Studies examining thrombocytopenia-associated mutations of these ABP show that all regulate proplatelet tip formation^3,7,8,31,32^. Reduced bending and bifurcation of the proplatelet shaft results in the release of fewer, larger thrombocytes, and in the case of NM-2A mutations, immature fractions are elevated^3,8,31^”^35^. Although proplatelets are not investigated, due to limited patient access, consistencies in thrombocyte properties between our study and these reports suggest proplatelet tip defects are likely in *ACTB*-TMS megakaryocytes. In support of this, we observe reduced levels and discontinuous staining of β-CYA and NM-2A in P5 (Fig. 4d) and P1 (Supplementary Fig. 6) thrombocytes compared with controls, indicating perturbation of this contractile system in *ACTB*-TMS peripheral blood thrombocytes.

Thus far, β-CYA disease-associated mutations have been biochemically characterized in a single study by our group, including the p.Glu364Lys mutation^36^, which is now incorporated into the *ACTB*-TMS cohort. We demonstrated that p.Glu364Lys mutant β-CYA can be recombinantly produced in the baculovirus/*Sf*9 expression system. Potent inhibition of DNase I activity showed that protein folding is minimally affected by this mutation. Experimentally verified functional features of the mutant protein such as reduced polymerization rate, impaired nucleotide exchange kinetics, and normal coupling between myosin binding and activation of ATPase activity were in good agreement with the results of MD-simulations. Accordingly, in this study we use *in silico* approaches to predict mutation mediated structural changes in β-CYA from P3 and P5 (β-CYA^P3^ and β-CYA^P5^). Both mutations perturb the structural integrity of SD1 (Fig. 5A and Supplementary Notes). The affected SD1 regions are within the interaction interfaces of NM-2 and α-actinin/filamin A (Fig. 5b and 5c).

**Figure 5.**
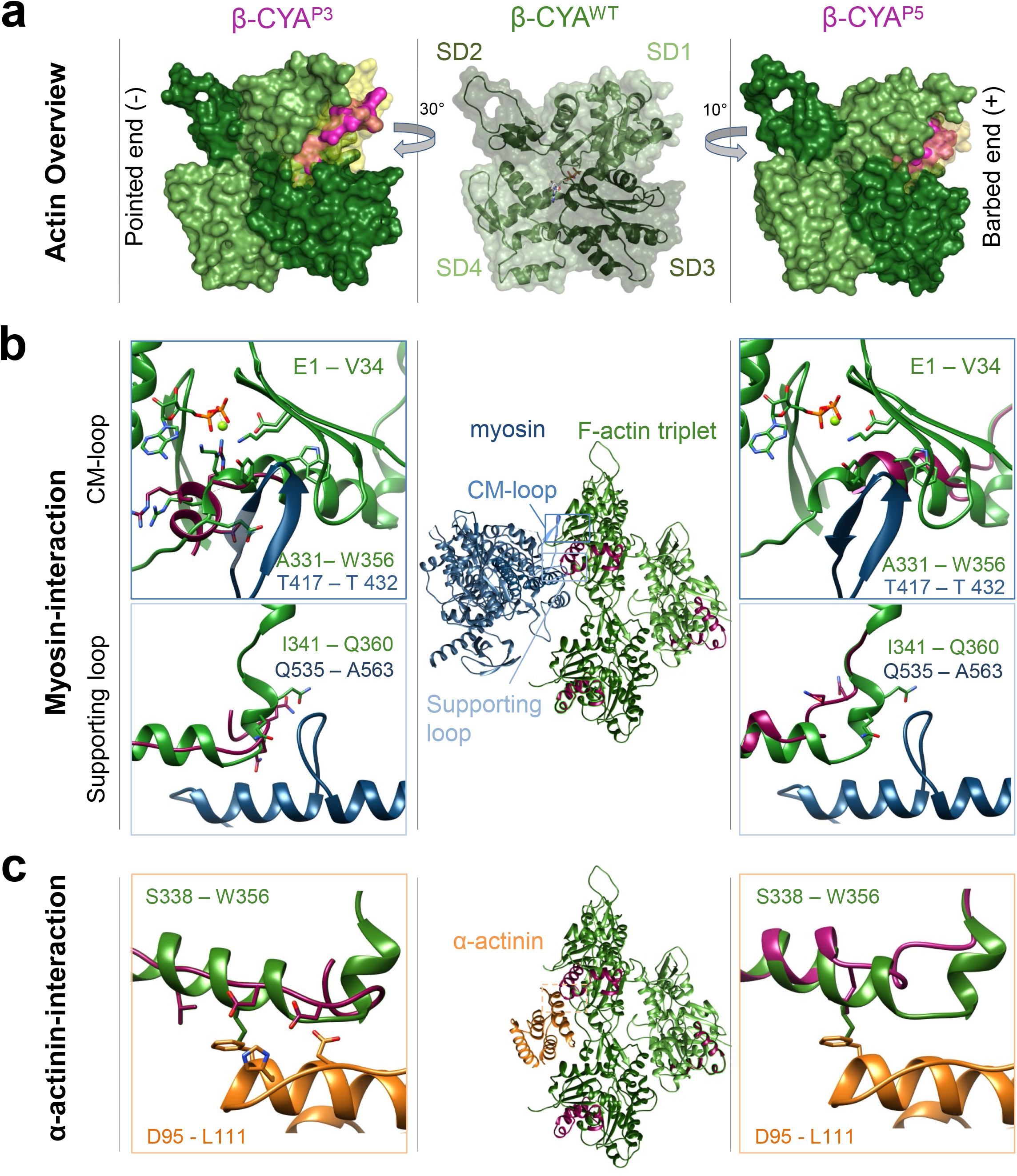
*In silico* modelling of P3- and P5-associated *ACTB* variants. (**a**) Overview of wildtype (WT) β-CYA (mid) in cartoon with space-fill overlay. β-CYA^P3^ and β-CYA^P5^ to the left and right respectively, with missing volumes indicated in yellow space-fill and affected residues modelled in magenta space-fill; (**b**) the binding interface NM-2 (blue, according to PDB 5JLH) on a β-CYA^WT^ F-actin triplet (mid, green with magenta C-terminal residues) is shown. Closeups of affected β-CYA^P3^ (left) and β-CYA^P5^ (right) residues are superimposed as magenta cartoon structures on the green β-CYA^WT^ cartoon structure. CM-loop interactions with mutant actins are shown in the top panels, whilst supporting loop interactions are modelled in the lower panels; (c) *In silico* modelling of the interaction interface of β-CYA^WT^ (green) with α-actinin (orange, according to PDB 3LUE, confirmed by docking) is shown (mid). Left and right show the interface with affected C-terminal residues for β-CYA^P3^ (left) and β-CYA^P5^ (right) respectively.

For NM-2, our model suggests significant perturbation for the cardiomyopathy (CM)-loop interaction with β-CYA^P3^ (Fig. 5b top). In the case of β-CYA^P5^, modelling and MD-simulations predict significant perturbations for interactions with both the CM-loop and supporting-loop, respectively^37,38^. For α-actinin/filamin A, hydrophobic interactions are conserved in both β-CYA^P3^ and β-CYA^P5^, however electrostatic interactions are introduced in β-CYA^P3^ (Fig. 5c). Whilst these models support altered interactions between mutant β-CYA and thrombocytopenia-associated ABP, we cannot exclude an effect of N-terminal α-SMA residues on their binding. However, the low level and variable incorporation of α-SMA within sub-nuclear bundles supports that mutant β-CYA is primarily responsible for the altered filament composition observed in *ACTB*-TMS cells. Further biochemical and structural studies are required to resolve the exact functional consequences of *ACTB*-TMS mutations at the molecular level.

In conclusion, this work describes a new clinical phenotype associated with mutations clustered in the 3’ region of *ACTB*, named *ACTB*-TMS. This is the first study to examine the pathogenesis of a human cytoplasmic actinopathy in patient derived cells. For two patients, we study the impact of these hypomorphic mutations at the clinical, cellular and molecular levels (summarized in Supplementary Table 7). Our results have identified a cytoskeletal phenotype unique to *ACTB*-TMS, whereby α-SMA is incorporated into mutant β-CYA bundles and a specific subset of ABP is recruited to these filaments. In dermal fibroblasts, these structures are linked with an increased contractile phenotype. We hypothesize that in megakaryocytes, they may limit the incorporation or capacity of essential ABP within proplatelets, compromising proplatelet tip deformation and thrombocyte maturation. Together, these data have resolved key components of a cytoskeletal filament population that both generates and transmits the forces required during the final stages of thrombopoiesis.

## Methods

### Patient recruitment

This study was approved by the TU Dresden institutional review board (EK127032017). Patients were recruited in four diagnostic laboratories after identification of likely pathogenic *ACTB* variants through whole exome sequencing (Agilent SureSelectXT kit, Agilent Technologies, Santa Clara, CA; P6), TruSight-One panel sequencing (Illumina, San Diego, CA; P5), and customized gene panels comprising 1800 rare disease genes enriched with a custom SureSelectXT Kit (Agilent, Santa Clara, CA; P1 and P2)) as well as 3089 genes associated with Mendelian diseases (P3 and P4, http://doro.charite.de/MENDEL_public/HPOv2_Gen_Panel.pdf). 100 or 150 nt single or paired-end sequencing was performed with a median target coverage of 80-fold either on an Illumina MiSeq, NextSeq500, or HiSeq Sequencing Systems. Alignment (mapping to GRCh37/hg19), variant identification (SNPs and indels), variant annotation and filtering was performed using the CLC Biomedical Genomics Workbench (Qiagen, Hilden, Germany)^39^. NextGENe^®^ software (SoftGenetics, LLC, State College, PA), Cartagenia Bench Lab NGS software (Agilent Technologies), or in-house software integrating open-source tools, such as BWA^40^, GATK^41^, FreeBayes toolkit, Jannovar^42^ and PhenIX^43^, were used.

### Primary fibroblast cell culture

Primary dermal fibroblasts were obtained from P3 and P5 following 3 mm cutaneous punch biopsies and cultured in BIO-AMF^™^-2 Medium (Biological Industries USA, Cromwell, CT, USA). For subculturing, primary fibroblasts were washed twice with 1× dPBS and detached at 37°C for 30 seconds with 1× Trypsin/EDTA (Gibco^®^; Thermo Scientific, Waltham, MA, US). The reaction was stopped with DMEM/F-12 medium (Gibco^®^; Thermo Scientific) containing 10% FCS (Gibco^®^; Thermo Scientific) and cells were pelleted by centrifugation at 500 *g* for 5 min. Cells were resuspended in BIO-AMF^tm^-2 medium, seeded onto Corning plasticware (Corning, NY, US) and maintained in BIO-AMF^tm^-2 medium at 37°C in the presence of 5% CO_2_. Cultures were continued for a maximum of 5 passages. For the growth curve assay, 3 × 10^4^ cells were seeded in 24-well plates and counted after 24 h, 48 h and 72 h following trypsinization. Culture images were obtained with a Nikon Eclipse TS100 microscope using 10×/0.25NA and 20×/0.40 NA objectives (Nikon, Minato, Tokyo, Japan).

### Flow cytometry

Patient-derived fibroblasts were cultured in BIO-AMF^™^-2 Medium in T75 flasks to a confluency of ~70%, washed with 10 ml 1X dPBS and detached using 2 ml Accutase^™^ (StemCell Technologies, Vancouver, Canada). After 5 minutes incubation at 37°C, the reaction was stopped by adding 2 ml medium. The cells were washed with dPBS containing 0.5% BSA and pelleted by centrifugation. The supernatant was discarded, and cells were fixed by dropwise adding of 1 ml 70% Ethanol cooled to - 20°C followed by gentle vortexing. Samples were stored at −20°C until the day of analysis and then pelleted upon adding of 6 ml 4°C dPBS. Two washing steps were carried out with 4 ml and 1 ml 4°C FACS-buffer (2 mM EDTA and 0,5% BSA in dPBS) and finally cells were resolved in 1 ml 2xUltraPure^™^ SSC (Invitrogen, Carlsbad, CA, USA). RNA-digestion was performed for 15 minutes at 37°C using 20 μg/ml RNase A (Qiagen) per sample. The samples were washed with 1 ml FACS-buffer and centrifuged for 10 minutes at 4°C and 300 g. For DNA staining, cells were resolved in 300 μl FACS-buffer containing 3 μM Propidium Iodite (Thermo Fisher) and incubated for 30 minutes in the dark at room temperature. All centrifugation steps were carried out at 260 g for 10 minutes if not otherwise mentioned. Samples were analyzed on a LSR Fortessa^™^ (BD Biosciences) of the Flow Cytometry Core Facility, a core facility of BIOTEC/CRTD at Technische Universität Dresden. Data were manually analysed using FlowJo analysis software (TreeStar, Ashland, OR, USA).

### Migration assay

CELLview^™^ 4 compartment glass bottom dishes (VWR International, Radnor, PA, US) were precoated with fibronectin/gelatin for 1 h at 37°C. Primary fibroblasts were seeded at a density of 2.5 × 10^3^ and allowed to attach overnight at 37°C in the presence of 5% CO_2_. Cells received fresh BIO-AMF^TM^-2 medium supplemented with 10 mM HEPES and dishes were mounted onto an inverted Nikon Eclipse Ti-E microscope system equipped with a 37°C incubator (Nikon). Phase contrast images were acquired at 10 min intervals for 8 h with a 10×/0.30NA objective. For analysis, cells were manually tracked with the MTrackJ plugin in ImageJ. Cells were excluded from the analysis if they collided with another cell or underwent division during the imaging period. The Dicty Tracking 1.4 software package was used to calculate cell trajectories, random migration speeds, directionality ratios and the mean square displacement of cells^44^.

### Cell lysis and western blot analysis

Cells grown to 90% confluency in 25 cm^2^ flasks were trypsinated as described above. Pelleted cells were resuspended, counted and washed twice in cold dPBS. Ice cold lysis buffer (20 mM Tris-HCl pH 8.0, 100 mM NaCl, 1 mM EDTA, 0.1% NP-40, complete mini protease inhibitor tablet and phosSTOP tablet (both from Roche Applied Science, Penzberg, Germany)) was added, giving a final concentration of 1 × 10^4^ cells per μl. Lysates were incubated for 30 min on ice, sonicated for 10 min in a water bath and vortexed. Total protein concentration was normalized with the Bradford Assay (BioRad, Hercules, CA, US) and readjusted using a Coomassie Protein Assay. Approximately 10–15 μg of total protein per sample was separated on 10% polyacrylamide gels by gel electrophoresis (35 mA, 45 min) and subsequently transferred onto 0.1 μm pore nitrocellulose membrane (14 V, 1.5 h). Transfer efficiency and protein loading were estimated by Ponceau S staining. Membranes were blocked for 1 h in 5% (w/v) skim milk in TBS-Tween20 (TBS-T) and incubated overnight in primary antibody diluted in blocking solution at 4°C (Supplementary Table 8). Following two TBS-T washes, membranes were incubated in secondary antibody in blocking solution for 1 h and washed thrice in TBS-T prior to developing. Signals were developed with the SuperSignal^TM^ West Femto Maximum Sensitivity Substrate (Thermo Scientific) and images were obtained with the ChemiDoc MP Imaging system using ImageLab software (Bio-Rad). All antibodies were tested twice with three separate lysates and the levels of β-CYA, γ-CYA, pan-actin and β-tubulin were assessed at least once in all lysates. Analysis of western blot signals was performed with Image J software (NIH, Bethesda, ML, US). Values were first adjusted to the total protein content, as determined by Ponceau S staining, and patient samples were normalized to their respective control.

### Primary fibroblast immunofluorescence microscopy

For immunofluorescence experiments, cells were seeded at a density of 2 × 10^4^ on 10 mm glass coverslips in 24-well plates and cultured for 48 h prior to collection. Cells were washed twice with pre-warmed DMEM/F-12 medium and fixed for 30 min with 1% PFA diluted in DMEM/F-12. Following two washes with 1× dPBS, cells were permeabilised with ice cold methanol for 5 min. Samples were washed twice with 1 × dPBS (Gibco^®^; Thermo Scientific) and blocked for 1 h at 23°C in 2% BSA/dPBS blocking solution prior to antibody labelling. Primary and secondary antibodies were diluted in blocking buffer, as shown in Supplementary Table 8, and were applied for 1 h and 30 min, respectively, with two intermittent blocking buffer washes. Samples were washed twice with 1 × dPBS, counterstained with DAPI for 5 min at RT, washed in 1× dPBS, rinsed with ddH_2_O and mounted in Prolong Gold anti-fade mounting medium (Thermo Scientific). Z-stack images (0.3 μm slices) were obtained with the Leica TCS SP8 Confocal Microscope (Leica Microsystems, Solms, Germany) using a 63× Oil/1.4NA objective. Image analysis was performed using Image J software. The cell attachment surface area was determined manually with ROI selection tools and the mean fluorescence intensity (MFI) of each channel was calculated from Sum Slice image projections using the same ROI. For overlap coefficient analysis, the ‘Just Another Colocalization Plugin’ was utilized^45^.

### Immunofluorescence microscopy of peripheral blood thrombocytes

For control/patient peripheral blood smears, slides were directly fixed in ice cold methanol at 4°C for 5 mins. All subsequent steps were as described above for primary fibroblast immunostaining. Micrographs with corresponding transmitted light images were collected at a single z-plane with a 60× Oil/1.35NA objective, 1024× 1024 pixel resolution and zoom factor of 3 using the Olympus FluoView 1000 (Olympus, Tokyo, Japan). Fluorescent micrographs were collected with the Leica TCS SP8 Confocal Microscope using a 63× Oil/1.4NA objective, 2048×2048 pixel resolution and 2.5 zoom factor. Z-stack slices had a spacing of 0.3 micron. Maximum intensity projections and fluorescence line profiles were obtained with Image J analysis software (NIH, Bethesda, ML, US). Whole-transcriptome sequencing (RNA-Seq)

For RNA-Seq, cells were seeded in 75cm^2^ flasks and harvested at ~70% confluence. RNA was extracted using the miRNeasy Mini Kit (Qiagen) according to the manufacturer’s instructions. On column DNA digestion was included to remove residual contaminating genomic DNA. All experiments were done in triplicate. For library preparations, TruSeq Stranded mRNA Library Prep Kit (Illumina) was used according to the manufacturer’s protocol, starting with 1 μg total RNA. All barcoded libraries were pooled and sequenced 2×75bp paired-end on an Illumina NextSeq500 platform to obtain a minimum of 10 Mio reads per sample.

Raw reads from Illumina sequencers were converted from bcl to fastq format using bcl2fastq (version v2.17.1.14) allowing for 1 barcode mismatch. Reads were trimmed for quality and sequence adapters using trimmomatic^46^. Trimming resulted in an average of 13.1–22.2Mio reads per sample. All reads were aligned against the phase II reference of the 1000 Genomes Project including decoy sequences d5 (ftp://ftp.1000genomes.ebi.ac.uk/vol1/ftp/technical/reference/phase2_reference_assembly_sequence/hs37d5.fa.gz) using STAR (v2.5.2b)^47^ in a 2-pass mapping mode: first, an index was created using the genome sequence and gene annotation (Gencode GRCh37.p13 comprehensive gene annotation), against which all reads are aligned. Second, all detected splice junctions of all samples are merged and used as guide for the second mapping step. Read counts of all annotated genes were extracted from the alignments using feature Counts method of the Rsubread package (v1.20.6)^48^ and genes with 0 counts for all samples were discarded. DESeq2 (v1.10.1)^49^ was used to find differentially expressed genes using standard parameters. The differentially expressed genes were filtered using the adjusted p-value <0.05. Significantly upregulated and downregulated genes were chosen based on different absolute log2 FC values.

Clustering was done with Euclidean distance and complete linkage using regularized-logarithm transformation (rlog) of TPM (Transcripts Per Kilobase Million) expression values. Heatmaps were plotted using the R package ComplexHeatmap^50^ and hclust of the R stats package (v3.4.2)^51^. Principal components analysis was done using the R stats package.

Pathway analysis was done using two different methods. First, Gene Ontology (GO) and KEGG enrichment of lists of differentially expressed genes with different absolute log2 FC cutoffs were calculated using DAVID Bioinformatics Resource^52^ via the R package RDAVIDWebService (v1.14.0)^53^ based on Ensembl IDs. The background set consisted of all genes passed to DESeq2. Second, the R package fgsea (v1.2.1)^54^ was used for a full gene set enrichment analysis. −log10(p-value) * signum(log2 FC) was used as rank function and 100,000 permutations for p-value calculation of pathway enrichments. KEGG pathways were plotted using the R package pathview (v 1.16.5)^55^.

A subset of 213 genes encoding for known actin binding proteins (based on ^56^) was analyzed further. As the actin cytoskeleton is abundant in mammalian cells, we rationalized that any proteins substantially influenced by the mutations and in turn influencing patient phenotype, should also be abundantly expressed. As such, genes were eliminated from the analysis when < 300 raw counts were observed in any of the samples analyzed. The remaining 124 ABP genes were sorted based on the log2 FC values calculated relative to the control. All genes upregulated and downregulated in both patients were then cross-checked on UniProt (http://www.uniprot.org/) and PubMed (https://www.ncbi.nlm.nih.gov/pubmed/) platforms for known associations with thrombocytopenia, as this is the clinical feature most consistent amongst the ACTB-TMS patients.

### *In silico* modeling of β-CYA^P3/P5^ and ABP interactions

Actin models of β-CYA^P3/P5^ were built with Yasara (version 17.8.15, YASARA Biosciences GmbH, Germany) based on PDB structure 5JLH^38^. For β-CYA^P3^, C-terminal residues after I330 were removed. The frameshift peptide was built via homology modeling using PDB structures as templates. The resulting peptide was placed manually into the model and was connected by the Yasara loop building algorithm. After energy minimization, an α-backbone constraint MD-simulation was performed with Desmond 11 MD package, version 2017-4, as distributed by Schrödinger^57^.The β-CYA^P5^ model was generated by deleting the residues spanning between Y337 and S344 and was subsequently closed with a VMD loop building algorithm. After energy minimization an α-backbone constraint MD-simulation was performed as described above. To confirm C-terminus stability, a 50 ns MD-simulation without constraints was performed with Desmond, using PDB 5JLH as a template. For the MD-simulations the OPLS3 force field was used in a minimized 10 Å orthorhombic water box with 0.15 M sodium chloride. The simulations were performed with a TIP3P water model including a 0. 03 ns quick relaxation. The simulation parameters were 300 K and 1 ATM pressure. MD-results were analyzed with VMD version 1.9.3^58^.To confirm the orientation of α-actinin binding on F-actin, the calponin-homology domain of PDB structure 3LUE^59^ was used in combination with the F-actin dimer from PDB structure 5JLH^38^. Frodock 2.0 software was used for docking^60^. To confirm stable binding, a 25 ns MD-simulation without constraints was performed with Desmond as described above. MD-simulation results were analyzed with VMD version 1.9.3^58^.

### Statistical analysis

Unless stated otherwise, GraphPad Prism 7.0 (GraphPad Software, San Diego, CA, US) was utilized for the graphical representation and statistical analysis. For statistical analysis, the Gaussian distribution of data was first assessed. Normally distributed data sets were analyzed with an ordinary one-way ANOVA followed by Holm-Sidak’s multiple comparisons test, whilst data that did not follow a Gaussian distribution were treated with the Kruskal-Wallis test followed by Dunn’s multiple comparisons test. In both cases, the mean ranks of all three conditions were compared with one another. In all cases, * = P<0.05, ** = P<0.01, *** = P<0.001 and **** = P<0.0001.

## Acknowledgments

We wish to thank the patients and their families, as well as the physicians and genetic counselors who referred them. We thank Christof Litschko, Institute for Biophysical Chemistry, Hannover Medical School for providing excel files for the evaluation of cell migration experiments and discussing the representation of cell migration data. Further, we acknowledge Dr. Rudolph Bauerfeind and the Laser Microscopy Core Facility at Hannover Medical School, and Dr. Andreas Pich and the Proteomics Core Facility at Hannover Medical School. We acknowledge Katharina Sarnow and Alexander Krüger for taking care of RNA extraction and RNA-Seq. Flow cytometry was performed with the support of the Flow Cytometry Core Facility, a core facility of BIOTEC/CRTD at TU Dresden. N.E. is a participant in the Berlin Institute of Health Charité Clinician Scientist Program, funded by the Charité - Universitätsmedizin Berlin and the Berlin Institute of Health. This work was financially supported by DFG grants MA 1081/22-1 and MA 1081/23-1 (D.J.M); Volkswagen Foundation grant VWZN3012 (D.J.M. and M.H.T); DFG grant DI 2170/3-1 (N.DD).

## Authors contributions

**S.L.L.** designed, performed and analyzed all cellular experiments. **N.DD.** and **N.E.** defined *ACTB*-TMS as a novel clinical entity. **R.K., N.E., M.J.L., M.J.F., J.A.L., K.B., T.M.N., E.S.**, and **A.R.** recruited patients, collected clinical data, and analyzed patient sequencing results. **M.C.F., R.H.** and **R.K.** analyzed blood and bone marrow smears and defined hematological phenotype of ACTB-TMS. **L.G.** performed flow cytometry. **K.S., K.G.**, and **B.K.** performed and analyzed whole-transcriptome sequencing. **P.Y.A.R.** and **D.J.M.** conceived structural studies. **P.Y.A.R.** performed *in silico* analysis. **M.H.T.** and **D.J.M.** supervised **P.Y.A.R.**, analyzed results, and contributed critical discussion. **C.C.** provided actin monoclonal antibodies and critical discussion for data interpretation. N.DD. coordinated the study and with **D.J.M.** co-sought funding. **S.L.L.** and **N.DD.** wrote the manuscript. All other authors critically analyzed and edited the manuscript.

## Additional Information

The authors declare no competing financial interests.

